# TOPII-targeting and MUS81 deficiency sensitise HER2-low tumour models to T-DXd

**DOI:** 10.64898/2026.08.04.742772

**Authors:** James Monypenny, Courtney Savage, Angela L. Caipa Garcia, Xiaolan Jiang, Gregory Weitsman, Marco Foiani, Tony Ng

## Abstract

In this study, we identify Topoisomerase II α (TopII α) targeting and MUS81 deficiency as sensitisers to trastuzumab deruxtecan (T-DXd), a HER2-targeting antibody–drug conjugate with a potent topoisomerase I poison payload that is of clinical importance in the treatment of HER2 expressing solid tumours in a range of cancer types. Using preclinical tumour spheroid models of breast and colorectal cancer, we show that TopII α targeting with doxorubicin sensitises HER2-low tumour cells to T-DXd, elevating cell cytotoxicity, DNA damage, and checkpoint pathway activation. T-DXd treatment, both as a single agent and in combination with doxorubicin, increases TopII α expression, highlighting this nuclear endonuclease as a potential candidate biomarker for T-DXd response in both the HER2-high and HER2-low setting. Using isogenic CRISPR models, we show that genetic loss of the MUS81 structure-specific endonuclease, a key processor of branched DNA structures and under-replicated DNA, sensitises HER2-low colorectal cancer cells to T-DXd. Given that reduced expression of MUS81 is closely related to metastasis and poor prognosis in colorectal carcinoma, our findings highlight the potential utility of these treatment combinations in a subset of colorectal cancer patients that present with HER2-positve/MUS81-low disease.

## INTRODUCTION

Amplified HER2 (*ERBB2*) expression is observed in 15-20% of breast cancers, where it promotes tumour survival and disease progression. Additionally, HER2 is amplified, over-expressed, and/or mutated in a variety of other solid tumours, including up to 30% of endometrial serous carcinomas, 15-20% gastro-oesophageal cancers, and 2-5% of lung and colorectal cancers. Oncogenic activation of HER2, a member of the EGFR family of protooncogene receptor tyrosine kinases, is commonly associated with amplified cell surface expression of the protein, making it an ideal target for antibody-drug conjugate (ADC) therapy.

ADCs represent a clinically significant advancement in monoclonal antibody-based therapy, offering great promise in oncology. Cytotoxic payloads conjugated to humanized monoclonal antibodies that target surface antigens enriched on tumour cells enable the delivery of drugs to the target site at concentrations that would otherwise be clinically intolerable by systemic administration of payload alone. ADCs therefore promote cytotoxic payload accumulation at the tumour site while reducing systemic payload exposure, with the aim of minimizing adverse drug reactions.

The rapid adoption and advancement in ADCs directed against HER2 has largely stemmed from the utilization of trastuzumab as the antibody backbone for these therapies. A wealth of data spanning two and a half decades related to the clinical utility and safety of trastuzumab has facilitated the accelerated development of HER2-directed ADCs that use this monoclonal antibody as a backbone (Baselga et al., 1999; Pegram et al., 1998). A well characterised and validated biological target, established diagnostics for patient selection, and regulatory approval for the carrier antibody have all contributed to the significant de-risking of HER2-directed ADC development, making it an attractive candidate for this promising area of cancer therapy.

T-DXd (trastuzumab deruxtecan, Enhertu) is the most recent HER2-directed ADC to receive clinical approval and represents a next-step advancement in ADC engineering (Ogitani et al., 2016). The parent antibody of T-DXd is trastuzumab, which is coupled to a potent topoisomerase I (TopI) inhibitor (exatecan derivative) via a cleavable tetrapeptide. The ADC has an impressive drug-to-antibody ratio (DAR; 8:1) and, compared with older generation ADCs, T-DXd exhibits a relatively homogenous DAR for improved pharmacokinetics. The TopI inhibitor payload is cell-permeable and, when liberated following cathepsin-mediated linker cleavage within the cellular lysosome, can diffuse locally within the tumour to poison ‘bystander’ cells that are not recipients of the ADC themselves. This ingenious approach is in-part designed to counter challenges posed by the heterogenous nature of HER2 expression often observed within tumours.

Within the UK and EU, T-DXd is employed in the treatment of early as well as advanced or metastatic HER2-positive breast cancers that have progressed on at least one prior anti-HER2 therapy, while in the US, T-DXd approval in the breast cancer setting has expanded to include HER2-low and ultra-low tumours (FDA, 2025). However, the clinical indications for which T-DXd is being adopted are progressively broadening to include a variety of solid tumour types that exhibit both high and low HER2 expression. Most notably, in 2024 and 2026, T-DXd received approval in the US and EU, respectively, for use in the treatment of any unresectable or metastatic HER2-positive (IHC 3+) solid tumours, landmark decisions that mark T-DXd as the first ADC to receive tumour-agnostic status, where clinical decisions governing therapy selection are primarily biomarker-driven (*Enhertu - Opinion on Variation to Marketing Authorisation (EMA)*, 2026; Research, 2024). These decisions were based on the findings from the DESTINY-PanTumor02, DESTINY-Lung01, and DESTINY-CRC02 Phase II trials that collectively revealed beneficial outcomes for T-DXd administered in HER2-positive disease over a broad range solid tumour types (Meric-Bernstam et al., 2024; Raghav et al., 2024; Smit et al., 2024).

The extended application of T-DXd to the HER2-low and HER2-ultra low setting within breast cancer, and to a broadening cancer landscape driven by a tumour agnostic clinical paradigm, warrants the identification of additional molecular markers that can inform on the likelihood of response, particularly in disease settings where HER2 may not be a key driver of tumor growth and progression. In this study, we identify Topoisomerase IIα (TopII α) and PARP as putative targets for enhancing T-DXd efficacy in HER2-low tumour spheroid models of breast and colorectal cancer (CRC). Additionally, our findings suggest that TopII α may have utility as a marker of response to T-DXd, whether alone or with combination treatments.

Through its unique, transient and tightly regulated endonucleolytic and strand-passage functions, TopII α processes DNA topological problems that are exacerbated following TopI trapping, including DNA supercoils, precatenanes and cruciform structures (Pommier et al., 2022; Wang, 2002). We demonstrate that targeting TopII α with low-concentration doxorubicin sensitises HER2-low cancer cells to the effects of T-DXd, elevating DNA damage, checkpoint pathway activation, G2M cell cycle arrest, and cell death.

Additionally, we show that T-DXd in combination with olaparib, a potent PARP1/2 inhibitor, dramatically elevated both DNA damage and the suppression of tumour cell growth, when compared with either agent alone. PARP plays a critical role in both the repair of single strand DNA (ssDNA) breaks and the removal of the toxic protein-DNA adducts that form in response to TopI poisoning. We show here that when used in isolation, olaparib has only a modest effect on DNA damage or the proliferation of tumour cells, but that it has a catastrophic effect on these processes in the T-DXd treatment setting. Notably, T-DXd, when used as a single agent or in combination with either doxorubicin or olaparib, results in elevated TopII α expression in both wild-type and MUS81-defficient cells, highlighting this enzyme as potential target and candidate biomarker for T-DXd response in both the HER2-high and HER2-low setting.

Using isogenic CRISPR models, we demonstrate that targeted depletion of the MUS81 structure-specific endonuclease sensitises HER2-low CRC tumour cells to T-DXd, elevating cell cytotoxicity, DNA damage, and checkpoint pathway activation. T-DXd combination therapy (doxorubicin and olaparib) further enhances the DNA damage and replication stress phenotype of CRC spheroids with a HER2-low/MUS81-defficient background. Given that reduced expression of MUS81 is closely related to metastasis and poor prognosis in colorectal carcinoma (Wu et al., 2011), our findings highlight the potential utility of these treatment combinations in a subset of CRC patients that present with HER2-positive/MUS81-low disease. This finding, coupled with the recent identification of heterozygous loss-of-function mutations in the *SLX4* gene in the T-DXd-resistant patient tumours, suggests that SLX4-MUS81-EME1 DNA damage response pathway alterations may provide potential mechanisms of escape from T-DXd payload cytotoxicity (Mosele et al., 2023). Based on our MUS81 data and these previously reported SLX4 findings, we hypothesise that attenuation of the late G2-stage functions of MUS81 may be implicated in T-DXd resistance in a manner that is uncoupled from the S-phase-dominant replication fork processing functions of this endonuclease.

## RESULTS

### The bystander antitumour effect enhances T-DXd efficacy in the 3D setting

To evaluate the relationship between ADC-dependent cytotoxicity and antigen expression, initial two-dimensional (2D) culture assays were performed using five BC cell lines that collectively display a broad spectrum of HER2 expression (Figure S1). Cell viability assays and western blotting demonstrate that, for both T-DM1 and T-DXd, ADC-dependent cytotoxicity is predominantly a function of cellular HER2 expression (Figure S1A). These data therefore confirm the assumption that HER2 expression is the major determinant of T-DM1 and T-DXd efficacy in 2D *in vitro* tumour models of breast cancer.

Immunofluorescence analysis of the DNA damage marker γH2AX in the HCC1954 HER2-high cell line revealed extensive induction of DNA damage following treatment with either ADC (Figure S1B), and viability data demonstrated both ADCs had a potent effect on the suppression of cellular growth. However, when these assays were performed within the three-dimensional (3D) setting using spheroid cultures, T-DXd was the more efficacious agent. Notably, immunofluorescence analysis of cryosectioned spheroids revealed that DNA damage was pervasive throughout the T-DXd treated spheroid, while it was more localised and most intense at the periphery for T-DM1 treated spheroids (Figure 1A). In addition to elevated DNA damage, suppression of spheroid growth, and cell accumulation in G2 were all more prominent for the T-DXd treatment group (Figure 1A, B, D-G). The greater overall DNA damage burden observed for T-DXd spheroids, as determined by western blotting analysis, was in contrast to that observed for the 2D culture setting, where T-DM1 was the more efficacious agent (Figure 1A; *cf.* bottom right blots).

**Figure 1:**
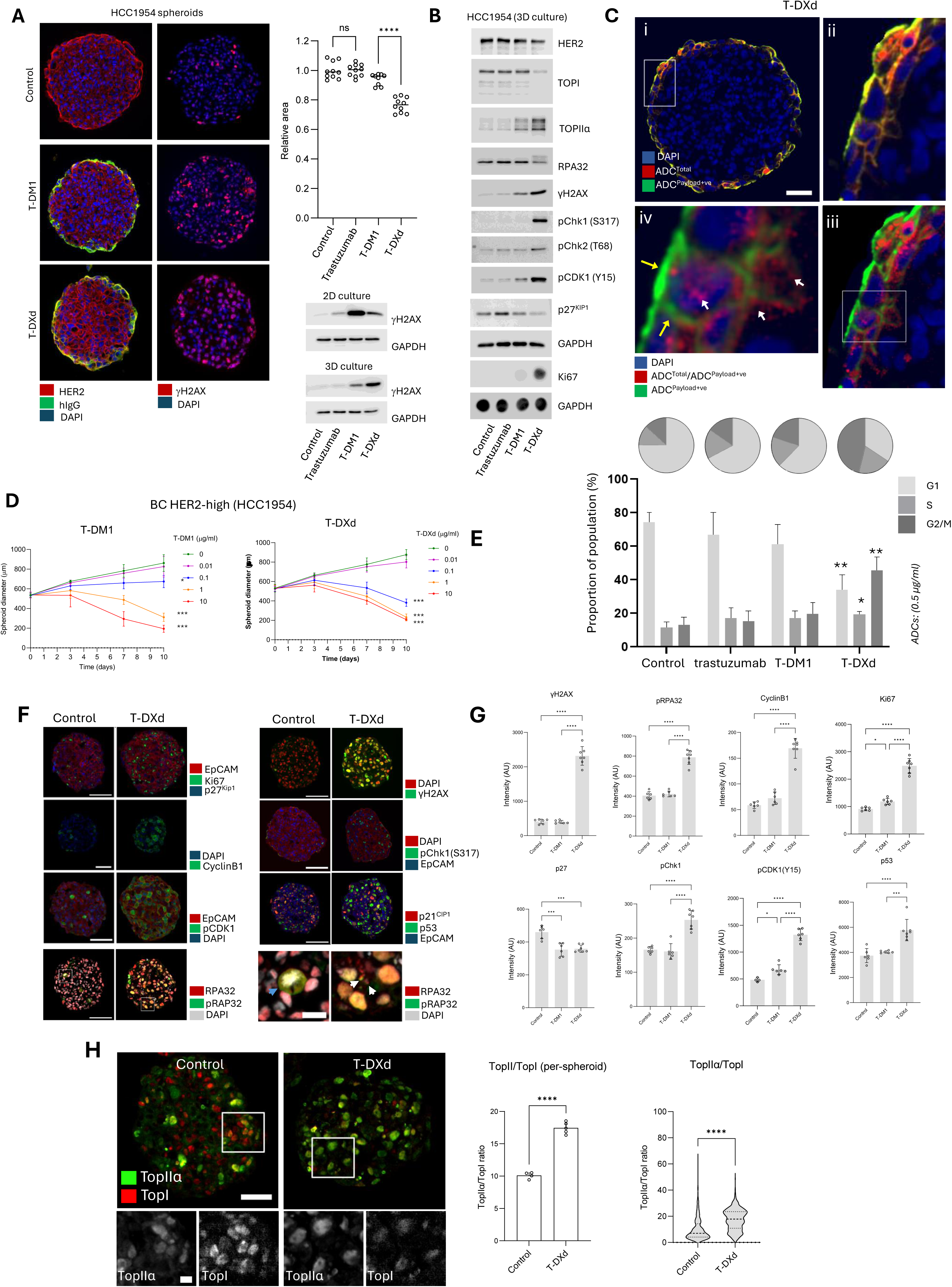
T-DXd demonstrates enhanced efficacy over T-DM1 in the 3D setting. (A) ADC studies performed using breast tumour spheroid models reveal enhanced T-DXd efficacy *vs.* T-DM1 in the three-dimensional culture setting. Immunofluorescence analysis (A; left image panel) of ADC distribution, HER2 expression and γH2AX induction in breast cancer spheroid cryosections. Membranous HER2 expression (A; left image column, red) is observed throughout the dense spheroid, with ADC engagement (green) detectable within only the first two-to-three cell layers of the perimeter. Analysis of γH2AX expression (right image column, red) shows T-DM1-associated DNA damage concentrated more within the cells of the spheroid periphery, while DNA damage associated with T-DXd is observed throughout the spheroid. Western blotting data (A; lower right panel) from ADC studies performed on two-dimensional *vs.* three-dimensional (spheroid) cultures, reveal a shift in T-DM1 and T-DXd efficacies between these different culture settings, as determined by γH2AX expression. Spheroid diameter analysis (top right graph) at D10 for the 0.1 µg/ml ADC treatment groups further demonstrates that T-DXd has a significantly greater suppressive effect on HCC1954 spheroid growth, when compared with T-DM1. (B) Western blotting and dot-blotting analysis of target protein expression in HCC1954 spheroids reveals that T-DXd promotes downregulation in TopI expression and a concomitant upregulation in TopII α expression. These changes are accompanied by a replication stress signature characterised by: (i) Chk1/2 phosphorylation, (ii) inhibitory CDK1(Y15) phosphorylation, (iii) RPA32 band-shifting (indicative of phosphorylation), (iv) γH2AX positivity and (iv) cell cycle stalling, as indicated by accumulation of S/G2-associated Ki67 expression and a concomitant reduction in G1-associated p27^Kip1^ expression. (C) Immunofluorescence analysis of ADC distribution using anti-payload and anti-backbone antibodies (upper panel) performed on T-DXd-treated HCC1954 spheroid sections. Upper images were processed to generate corresponding image maps (lower panel) of the per-pixel relative proportions of processed (hIgG+ve/payload+ve) *vs.* unprocessed (payload+ve) within the cells of the spheroid. Mapping of ADC processing reveals the highly efficient liberation of payload upon ADC internalisation (as demonstrated by the high hIgG+ve/payload+ve signal). (D) Spheroid growth assays demonstrate the enhanced efficacy of T-DXd over T-DM1 in the three- dimensional culture setting (N = 3 repeats). (E) Flow cytometry analysis of cell cycle phase reveals T-DXd- dependent HCC1954 cell trapping within both the S and G2/M phases (upper pie charts, representative experiment; lower bar graph, statistical comparisons from N = 3 repeats). (F) Immunocytochemistry analysis reveals the spatial expression of cell cycle, replication stress and DDR markers associated with the T-DXd-associated biochemical phenotype. Representative immunofluorescence images of Ki67, CyclinB1, inhibitory CDK1 phosphorylation, pChk1, γH2AX, and pRPA32 expression in HCC1954 spheroid sections from control and T-DXd treatment groups. Note that pRPA32 is abundant and associated with pre- mitotic cells, which are largely absent in T-DXd-treated cultures (see enlarged images of regions of interest in the bottom panel; blue arrowhead: mitotic cell in control group). Enlarged images also reveal the presence of micronuclei in T-DXd-treated cells (white arrowheads). (G) Statistical analysis of image data, showing per-spheroid mean immunofluorescence intensities for indicated DNA damage and replication stress markers (N≥4 independent spheroids per treatment group) (H) Representative image of dual TopI/TopII α immunofluorescence staining (left image panel) in control and T-DXd-treated HCC1954 BC spheroids, showing TopI downregulation and a concomitant upregulation in TopII α expression following T- DXd treatment. Statistical analysis of whole spheroid (middle graph) and per-nuceli TopII α/TopI intensity ratios performed on immunofluorescence data reveal a statistically significant increases in the mean TopII α/TopI ratio following T-DXd treatment.

Co-labelling of the ADC within these spheroids revealed it to be primarily restricted to the outmost cellular layers of these structures (Figure 1A), suggesting bystander effect as the mechanism for the extended cytotoxic reach observed in these 3D cultures following T-DXd treatment. To investigate this further, image analysis was performed on spheroid sections using antibodies targeting both the ADC antibody backbone and the ADC-bound payload. This analysis revealed the highly efficient processing of the internalised drug within the outermost cell layers of the spheroid periphery (Figure 1C), as determined by ADC backbone-high, but payload-low signal (liberated payload is lost post-fixation and permeabilization). This analysis suggests highly efficient payload processing at the spheroid periphery is sufficient to poison cells deeper within the core due to bystander reach.

### Enhanced TopII α expression is a marker of T-DXd response

In addition to a reduction in viability, elevated expression of replication stress markers was a key hallmark of T-DXd-depended cytotoxicity (Figure 1B, F & G). Activation-associated phosphorylation of the checkpoint proteins Chk1/Chk2, the inhibitory phosphorylation of CDK1 at Y15, and ATR activation-associated phosphorylation of RPA32 on S4/S8 were all elevated following exposure of HCC1954 spheroids to T-DXd (Figure 1B, F & G). Notably, elevated p53 expression was also observed in cells throughout these spheroids.

Reduced TopI expression was also observed in HCC1954 cells in the T-DXd-challenged spheroid (Figure 1B & H), which was a possible consequence of the proteolytic trimming and subsequent TDP1-dependent removal of persistent poisoned TopI-DNA covalent complexes (TopIcc) (Sun et al., 2020). Interestingly, this TopI loss was mirrored by a concomitant increase in TopII α expression (Figure 1B & H), with the ratio in TopII/TopI expression providing a sensitive metric for drug response in HCC1954 cells at both the spheroid and cellular level (Figure 1H, left and right chars, respectively). TopII α is a cell-cycle regulated gene, showing peak expression during S, G2 and M phases of the cell cycle (Goswami et al., 1996; Pommier et al., 2022). Similarly, the expression of the cell cycle regulated genes Ki67 and Cyclin B1, the latter of which parallels TopII α expression, were also elevated in T-DXd treated spheroids. This somewhat paradoxical observation (apparent increase in proliferation index; figure 1G, top right graphs) was a consequence of an enriched G2 cell population, due to replication-stress induced checkpoint activation and cell arrest.

The three different spheroid models employed in this study (HCC1954, T47D, DLD1) all display a similar and distinct anatomy that consists of a peripheral zone of proliferation (Ki67^high^/p27^low^) that transitions to an underlying zone of quiescent cells (Ki67^low^/ p27^high^) and then, in larger spheroids, a central necrotic core (this is most apparent in the fast-growing DLD1 line, Figure S1C). TopI poisoning disrupts this anatomy, greatly enlarging the Ki67^high^/p27^low^ cell population. This is a likely consequence of incomplete cell cycle progression as cells become arrested and trapped within the S/G2/M phases of the cell cycle (Figure 1E & F), resulting in the accumulation of an expanding zone of Ki67^+ve^ but replication defective cells. Consequently, in addition to the contribution of a bystander effect, this accumulation of trapped, replication deficient cells likely also contributes significantly to the expansive elevated expression of γH2AX, proliferation markers, and checkpoint pathway proteins throughout the T-DXd challenged spheroid.

### TOPII targeting sensitise HER2-low tumour models to T-DXd

TopII α is known to share functional overlap with TopI. The observed elevation in TopII α expression in T-DXd treated cells therefore presents the possibility that TopII α may partially compensate for TopI functional loss or that it may process aberrant DNA topologies arising from TopIcc-associated genotoxicity, to counter the effects of T-DXd. To address this question, we explored the effects of T-DXd alone or in combination with the TopII α inhibitor doxorubicin in HER2-low tumour models, given the already high toxicity observed for T-DXd in the HER2-high HCC1954 cell line. The T47D breast and DLD1 colorectal cancer models were therefore selected for subsequent studies, given their established low HER2 expression and their excellent spheroid formation and growth characteristics.

In agreement with 2D cell culture assay data (Figure S1A), T47D spheroids were less sensitive to T-DM1 and T-DXd, when compared with the HCC1954 tumour model (Figure 2A & B vs. 1D & E), a likely consequence of their low level of HER2 target antigen expression (Figure S1A).

**Figure 2:**
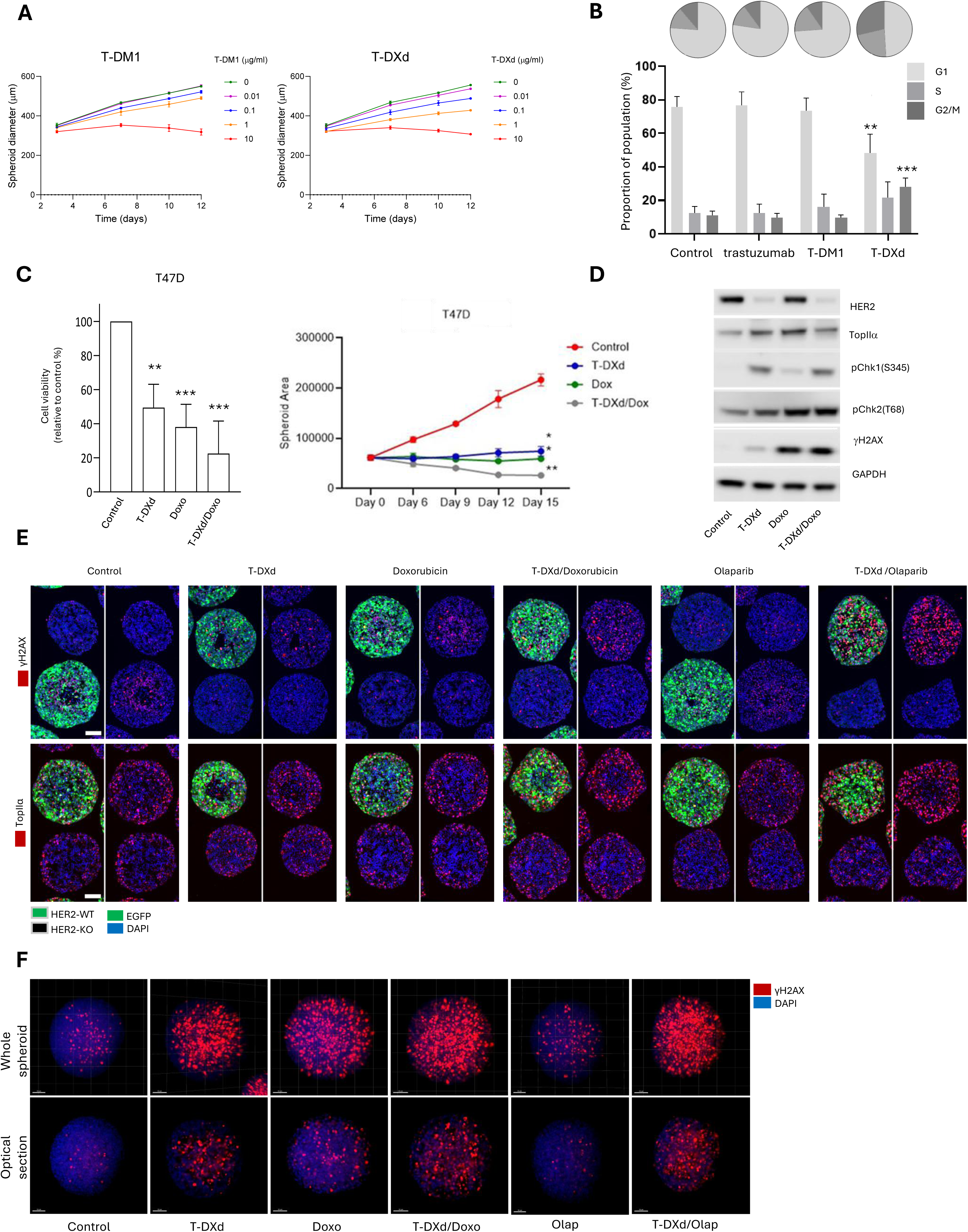
TOPII targeting sensitise HER2-low tumour models to T-DXd. (A) Spheroid growth assays and (B) cell cycle analysis demonstrating the inhibitory effects of T-DM1 and T- DXd on the HER2-low T47D BC cell line. Cell cycle analysis performed at D3 post treatment (upper pie charts, representative experiment; lower bar graph, statistical comparisons from N = 3 repeats, trastuzumab/ADCs used at 0.5 μg/ml). (C) Cell titer glow-based viability (left-hand graph) and spheroid growth assays (right-hand plot) demonstrating the effects of T-DXd and doxorubicin, either as single agents or in combination, on T47D HER2-low BC spheroids N=3 repeats; **, p < 0.01, *** p < 0.001). (D) Western blot demonstrating the effects of the single agents and combination therapy on DNA damage, checkpoint pathway activation, and TopII α expression in day-three treated T47D HER2-low BC spheroids (representative of N=2 blots). (E) Comparative immunofluorescence analysis of γH2AX and TopII α within a mixed population of HER2-WT and HER2-KO T47D spheroids, following challenge with the indicated single agent and combination therapies. HER2-WT spheroids are identified by the co-expression of EGFP, while DAPI labelling enables identification of both spheroid populations. This experiment setup enables direct comparisons between test HER2-WT and control HER2-KO spheroids within a given treatment group, as well as across treatment groups, all combined together on the same imaged slide (See figure S3B C C for methodology overview). (F) Light-sheet imaging of cleared whole-mount T47D breast cancer spheroids labelled for DAPI (blue) and γH2AX (red). The images of the upper row show a full projection of the three- dimensional spheroid while the images of the lower row show a single corresponding optical section, for comparison with cryosection data in the upper panel (γH2AX) of E.

However, despite the extremely low level of HER2 expression in the T47D cell line, spheroid growth assays revealed a none-the-less relatively robust response to T-DXd (Figure 2A, Figure S2A & B). And similarly to HCC1954, T-DXd demonstrated greater efficacy than T-DM1, both in terms of spheroid growth suppression and the accumulation of cell populations within the S and G2/M phases of the cell cycle (Figure 2A & B). Given these observations and the known low level of HER2 expression in these cells, it was important to confirm that the cytotoxic effects of the treatment were attributable to antigen-mediated uptake of the ADC and not due to off-target effects associated with passive uptake of drug over a prolonged treatment period. A syngeneic T47D HER2-knockout CRISPR model was therefore generated to discriminate between the on-target and off-target effects of ADC under our experimental conditions. Cytotoxicity assays conducted using this model enabled the unequivocal confirmation of antigen-mediated (on-target) uptake and delivery of T-DXd payload at concentrations of 1 µg/ml and below for an exposure period of up to two weeks (Figure S3A & B). At concentrations above 1 µg/ml, off-target mechanisms were found to significantly contribute to suppression of spheroid growth. Therefore, all subsequent studies employed T-DXd at a concentration of 0.5 μg/ml, which was one-half of the maximum safe on-target concentration established from CRISPR assays.

To evaluate whether TopII-targeting may sensitise T47D spheroids to T-DXd and thereby extend the ADC’s efficacy in the HER2-low setting, spheroids were exposed to a combination of T-DXd and doxorubicin, a TopII poison. Cytotoxicity and spheroid growth assays revealed greater suppression of spheroid growth under the combination therapy setting, when compared with either drug alone (Figure 2C). Biochemical analysis of checkpoint pathway activation revealed that, as single agents, T-DXd robustly activates Chk1, while doxorubicin exclusively activates Chk2 (Figure 2D), suggesting that the robust activation of both arms of the checkpoint pathway may in part account for the enhanced suppression of spheroid growth and increased cytotoxicity observed under the combination therapy setting.

### T-DXd combination therapy elevates global DNA damage and enhances TopII α expression in HER2-low breast cancer spheroids

To further evaluate the effects of these treatments on DNA damage and TopII α marker expression *in situ*, we developed a sample processing and imaging workflow that enables spheroid cryosections from multiple treatment groups to be imaged together on the same slide, with each treatment group containing populations of syngeneic HER2-wildtype (HER2-WT) and HER2-knockout (HER2-KO) spheroids, the latter serving as an internal control for baseline marker expression and potential off-target cytotoxicities (Figure S3B-D). For all imaging assays, we included olaparib as a positive control for the combination therapy setting, given its known strong interaction with TopI poisons but otherwise low cytotoxicity when used as a single agent. Using this workflow, immunofluorescence analysis revealed that T-DXd/doxorubicin and, in particular, T-DXd/olaparib combination therapies resulted in a more robust elevation in γH2AX and TopII α marker expression, when compared with spheroids treated with single agents (Figure 2E). In addition to our spheroid sectioning-based immunofluorescence pipeline, we also employed light-sheet microscopy in the analysis of whole-mount specimens, to evaluate the effects of combination therapies *vs.* single agents on DNA damage throughout the entire spheroid. While analysis of γH2AX expression in spheroid sections initially suggested that the effects of T-DXd alone or T-DXd/doxorubicin combination therapy on elevated DNA damage were relatively modest, whole-spheroid imaging of γH2AX expression revealed more completely the extent of this damage, which could be observed from the spheroid periphery deep into the core (Figure 2F). Importantly, individual optical sections from light-sheet image stacks revealed a pattern of γH2AX expression that was similar to that observed for the sectioning-based image workflow, indicating good concordance among the assays. Taken together, these findings demonstrate that the enhanced cytotoxic effects of T-DXd/doxorubicin and T-DXd/olaparib combination therapies towards HER2-low breast cancer cells are associated with extensive DNA damage, elevated TopII α expression, and robust Chk1/Chk2 pathway activation.

### MUS81 loss sensitises HER2-low CRC spheroids to T-DXd

To extend our analysis of T-DXd combination therapy efficacy beyond the breast cancer setting, we next performed studies using the DLD1 cell line which is classified as microsatellite-unstable, as well as an established HER2-low model of colorectal cancer that demonstrates excellent spheroid growth characteristics (van Wietmarschen et al., 2020). These microsatellite unstable cells likely contain TA repeats in the genome which in turn are particularly prone to form cruciform structures targeted by MUS81. In addition to testing T-DXd in combination with doxorubicin and olaparib within the CRC setting, we also sought to evaluate the contribution of the MUS81 endonuclease to the T-DXd response given the previously reported association between this key replication stress and DNA damage response factor and CRC prognosis and metastasis. CRIPSR-mediated targeted deletion of *MUS81* was performed in DLD1 cells and three expanded MUS81-negative clones then pooled to generate a MUS81-KO syngeneic line (Figure S4, left-hand blot). HER2 and TopI levels in the MUS81-KO line were compared to those of the MUS81-WT line to ensure matched expression of T-DXd antigen and payload targets (Figure S4, right-hand blot).

Spheroid growth and cell cytotoxicity assays were performed to evaluate the effects of T-DXd, either as a single agent or combination with doxorubicin, in both the DLD1 MUS81-WT and MUS81-KO background (Figure 3A & B). These assays revealed that, in the MUS81-WT setting, single-agent T-DXd and doxorubicin had a similarly suppressive effect on DLD1 spheroid growth that appeared cytostatic over the two-week experimental period (Figure 3A, top graph). T-DXd/doxorubicin combination therapy, however, had a more suppressive effect than either agent alone, resulting in a gradual reduction in spheroid size over time. In the MUS81-KO setting, T-DXd had a markedly more suppressive effect of spheroid growth, when compared with doxorubicin, resulting in a sustained decrease in spheroid size over the course of the experiment. T-DXd/doxorubicin combination therapy had little benefit over T-DXd alone in this setting, given the already highly growth-suppressive and cytotoxic effects of the ADC in the MUS81-KO background (Figure 3A, bottom graph; Figure 3B). MUS81 loss also promoted sensitivity to olaparib in DLD1 spheroids, while T-DXd/olaparib combination therapy was equally and highly efficacious in both a MUS81-WT and MUS81-KO background (Figure 3B).

**Figure 3:**
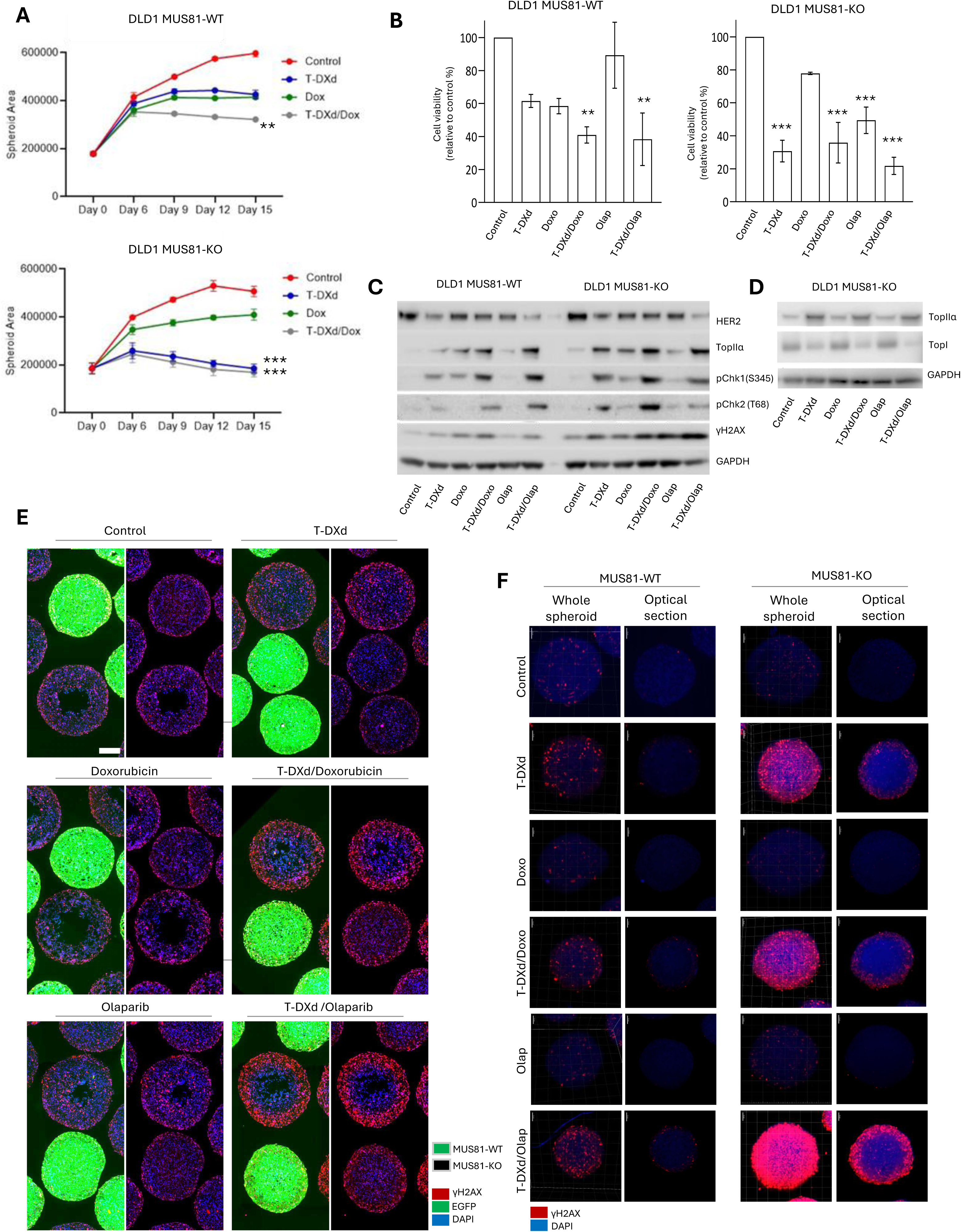
MUS81 loss sensitises HER2-low CRC spheroids to T-DXd and T-DXd/doxorubicin combination therapy. (A) DLD1 MUS81-WT (upper) and MUS81-KO (lower) spheroid growth assays showing the change in mean spheroid diameter over time for the indicated treatment groups. Statistical comparisons shown were performed on mean spheroid diameter measurements obtained at day 12 (N=3) (B) Cell viability determined by cell titer glow assay for DLD1 WT (left-hand graph) and MUS81-KO (right-hand graph) spheroids subjected to the indicated treatments and then processed at day 12 (N=3) (C) Western blot demonstrating the sensitising effects of MUS81 loss on DNA damage and checkpoint pathway activation in DLD1 spheroids challenged with T-DXd alone or in combination with doxorubicin or olaparib (representative on N=3 repeats). (D) Western blot showing TopII α upregulation and concomitant TopI downregulation in MUS81-KO DLD1 spheroids challenged with the indicated therapies (representative of N=2 repeats). (E) Comparative immunofluorescence analysis of the DNA damage marker γH2AX within a mixed population of MUS81-WT and MUS81-KO DLD1 spheroids, following challenge with the indicated single agent and combination therapies. MUS81-WT spheroids are distinguishable by the expression of EGFP. (F) Light-sheet imaging of cleared whole-mount MUS81-WT (left panel) and MUS81-KO (right panel) DLD1 spheroids labelled for DAPI (blue) and γH2AX (red). For each panel, the images of the left-hand column show a full projection of the three-dimensional spheroid while the images of the right-hand column show a single corresponding optical section, for comparison with cryosection data in E.

Analysis of protein lysates from day-three treated spheroids (Figure 3C) revealed that the T-DXd combination therapies had the greatest effect on Chk1/Chk2 pathway activation in the MUS81-WT setting, while T-DXd and T-DXd/doxorubicin combination therapy resulted in similar and robust Chk1/Chk2 activation in the MUS81-KO setting. T-DXd, either alone or in combination with doxorubicin or olaparib, resulted in the greatest increase in TopII α expression in the MUS81-KO background, while this phenotype was most pronounced for the combination therapies in the MUS81-WT background. In the MUS81-KO background, the elevated TopII α expression associated with the T-DXd treatment groups was mirrored by a concomitant reduction in TopI expression. This was similar to that observed in the HER2-high HCC1954 breast cancer cell line and suggests that under conditions of both MUS81 loss and T-DXd challenge, degradation of poisoned TopI complexes outpaces *de novo* protein synthesis (Figure 3D). Double-strand DNA breaks, as determined by γH2AX expression, was greater for all given treatment groups of the MUS81-KO setting, when compared with the MUS81-WT setting.

An *in situ* analysis of DNA damage in DLD1 spheroids using our cryosectioning and light-sheet microscopy-based immunofluorescence pipelines revealed extensive γH2AX expression throughout the peripheral zone of the MUS81-KO spheroid (Figure 3E & F). DNA damage was most extensive for the T-DXd/doxorubicin and T-DXd/olaparib treatment groups and was far greater for MUS81-KO setting than for MUS81-WT setting. Again, good concordance was observed in the γH2AX phenotype between cryosections and optical sections for the two different imaging assays.

### Persistent DNA damage, elevated TopII α expression and G2/M arrest are markers of T-DXd response in tumour spheroids chimeric for MUS81 expression

In addition to the application of light-sheet microscopy, a chimeric spheroid experimental model was developed to enable the direct comparison of MUS81-WT and MUS81-KO DLD1 tumour cells within the same spheroid, to address some of the technical challenges posed when making inter-spheroid comparisons of marker expression within sectioned specimens (e.g., alterations in marker expression as a consequence of section depth). The image processing pipeline maps and extracts the MUS81-WT (identifiable by EGFP expression) and MUS81-KO cell populations on a per-spheroid basis. Subsequent nuclear masking is performed to extract mean fluorescence intensity values for the markers γH2AX and TopII α to facilitate a statistical evaluation of inter-population differences in marker expression (Figure 4A C B).

**Figure 4:**
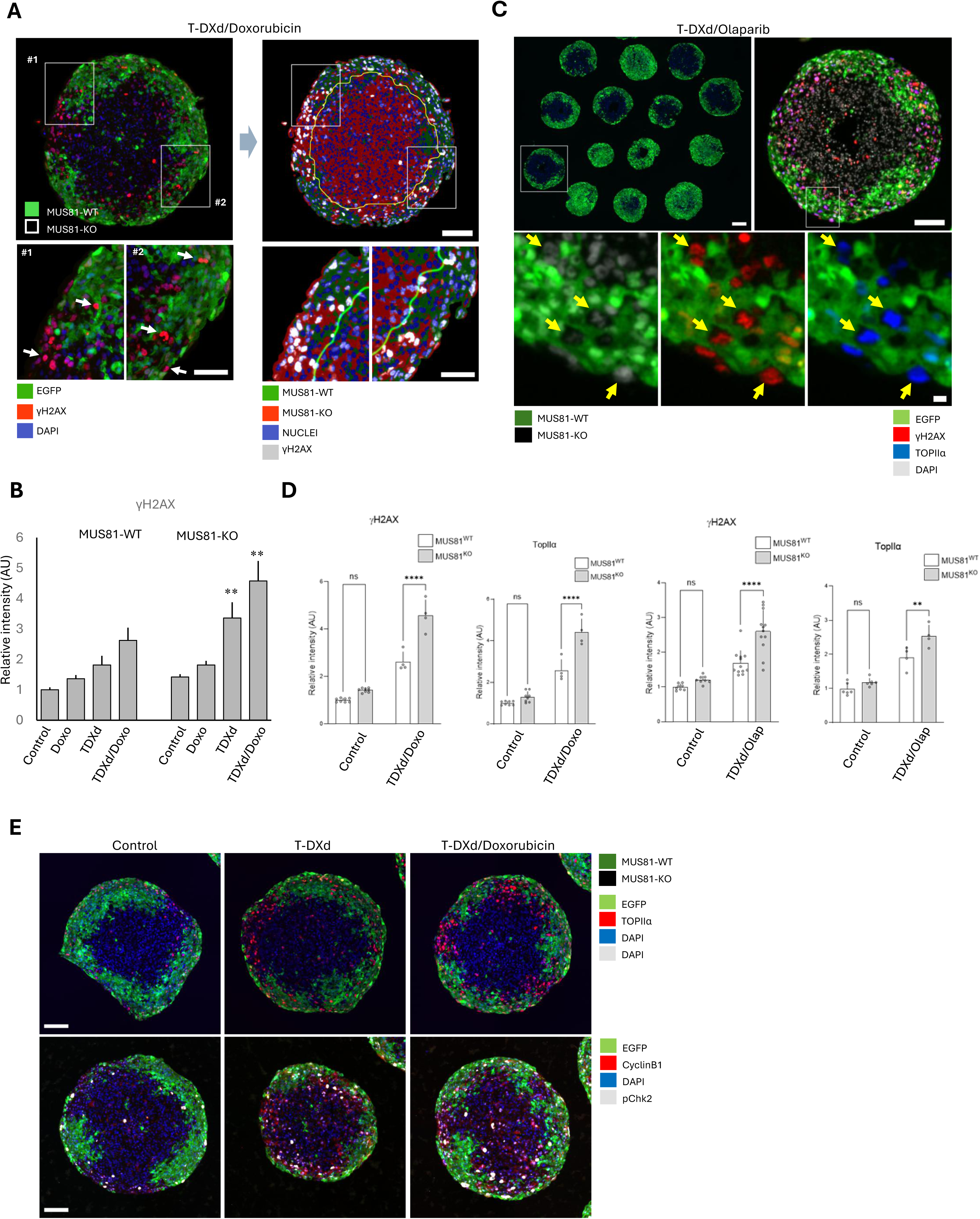
Engineered zones of MUS81-deficiency within the tumour spheroid highlight DNA damage and TopII α marker utility in the T-DXd combination therapy setting. A chimeric spheroid experimental model was developed to enable the direct comparison of MUS81- WT and MUS81-KO DLD1 tumour cells within the same spheroid, to address some of the technical challenges posed when making inter-spheroid comparisons of marker expression within sectioned specimens. (A) Immunofluorescence analysis of chimeric MUS81-WT/ MUS81-KO DLD1 spheroid sections revealed elevated and extensive DNA damage within the MUS81-KO population, confirming the sensitising effects of MUS81-loss on T-DXd/doxorubicin combination treatment and extending this observation to specimens of mixed MUS81 expression. Arrows highlight examples of intense γH2AX nuclear staining in MUS81-KO cells (EGFP-negative). Right hand panel: Representative example of the image processing pipeline used to map and extract the MUS81-WT and MUS81-KO cell populations on a per-spheroid basis, to enable subsequent nuclear masking and extraction of per-population marker intensities (γH2AX, TopII α). (B) Bar graph summarising quantitative analysis of γH2AX intensities within the nuclei of MUS81-WT and MUS81-KO cell populations on a per-spheroid basis for the indicated treatment groups. An asterisk denotes a significant difference in intensity of expression between the MUS81-WT and MUS81-KO population for the indicated treatment group (N≥4 independent spheroids per treatment group). (C) Immunofluorescence analysis of spheroid sections of chimeric MUS81-WT/ MUS81-KO DLD1 spheroids exposed to T-DXd/olaparib combination therapy, demonstrating the efficacy of this combination treatment strategy in the MUS81-deficient setting. (D) Bar graph summarising per-spheroid γH2AX and TopII α mean nuclear intensities extracted from image data for the MUS81-WT and MUS81-KO cell populations of the control and T-DXd combination therapy treatment groups (N≥4 independent spheroids per treatment group). (E) Immunofluorescence analysis of cryosections of chimeric MUS81-WT/MUS81-KO DLD1 spheroids stained for TopII α (top panel) and CyclinB1 and pChk2 (bottom panel) at day three-post indicated treatments.

Image analysis of chimeric DLD1 syngeneic MUS81-WT/ MUS81-KO spheroids revealed treatment dependent elevation of DNA damage and TopII α expression that was significantly enriched within the MUS81-deficient cell population (Figure 4C & D). T-DXd combination therapies had the most striking effects on these phenotypes. The elevated expression of γH2AX and TopII α was also associated with a robust induction of pChk2 activation and a cytoplasmic accumulation of CyclinB1 (Figure 4E), suggesting that T-DXd and T-DXd combination therapies are strong drivers of G2/M checkpoint arrest in the MUS81-defficient setting.

## DISCUSSION

In this study, we employ tumour spheroid cultures and a range of viability, biochemical, and imaging assays to explore effective therapeutic combinations and potential markers of response for the T-DXd treatment setting, with a focus on models of HER2-low disease. We show that T-DXd combination treatment with doxorubicin or olaparib enhances response in HER2-low spheroid models of breast and colorectal cancer. We show that T-DXd treatment promotes arrest in the S and G2 phases of the cell cycle, and elevates TopII α expression, highlighting the potential utility of this enzyme as a marker of treatment response. We further show that MUS81 loss sensitizes HER2-low colorectal cancer spheroids to T-DXd, either as a single agent or in combination with these therapies. There is a strong rational for exploring the interaction between TopI poisons, TopII inhibition, and MUS81 deficiency, both mechanistically and clinically.

During S phase, TopI relieves replication and transcription-associated torsional stress in the DNA double helix by inducing a transient nick in the phosphodiester backbone of one of the duplexed strands. This promotes a single rotation along the axis of the double helix that dissipates torsional stress. Subsequent re-ligation of the phosphodiester backbone completes the enzymatic cycle. During this cycle, TopI becomes transiently covalently coupled to DNA, forming a structure known as the TopI covalent complex (TopIcc). DXd, the payload of T-DXd, as well as other clinically approved TopI poisons, specifically target TopIcc, trapping the enzyme-DNA complex (Pommier, 2006). This has three consequences that are deleterious for replication; the persistence of a single strand break in the DNA backbone, the generation of a bulky protein-DNA adduct, and the unproductive dissipation of torsional stress within the double helix. If unresolved, trapped TopIcc can promote replication run-off (and thereby a genotoxic one-ended double-strand break), replication fork stalling and/or reversal, and ultimately fork collapse. Additionally, TopI poisoning impacts on transcriptional machinery, increasing the genomic R loop burden as unresolved negative torsional stress increases the favorability of DNA-RNA hybridization (Duardo et al., 2024; Pommier et al., 2022). R loops stall transcription and present barriers to replisome progression, promoting transcription-replication conflicts during S phase (Saha & Pommier, 2023). Collectively, these processes drive DNA damage and elevate replication stress, and would be predicated to increase the burden of aberrant DNA topology and under-replicated DNA of cells that transit into G2. In the current study, our flow cytometry and imaging data demonstrate that T-DXd promotes an accumulation of cells within the S and G2 phases of the cell cycle (Figures 1E-G, 2B & S2B, 4E). In the context of TopI poisoning, cells that transit into G2, will likely do so with an increased burden of under replicated DNA and aberrant DNA topology, both of which must be resolved to enable faithful cell division (Bertolin et al., 2020; Pommier et al., 2022). During G2, TopII α- and MUS81-dependent pathways cooperate to resolve this defective state.

TopI and TopII inhibitors are known to perturb different temporal windows of genome duplication and chromosome resolution, with TopI poisons primarily inducing the inter-S phase checkpoint, and TopII poisons driving G2/M arrest (Xiao et al., 2003). TopI and TopII also share overlapping substrates and likely exhibit a degree of functional redundancy (Bermejo et al., 2007; Pommier et al., 2022). Consequently, T-DXd-dependent TopI poisoning would be predicted to increase reliance on TopII α for the resolution of aberrant DNA topology, including persistent supercoils and precatenanes during S phase. In our study, we identify TopII α upregulation as a potential marker of T-DXd response, both in the HER2-high (Figure 1B & H) and HER2-low (Figures 2D, S2A, & 3C) setting. Given that TopII α is a cell cycle regulated gene (Goswami et al., 1996; Pommier et al., 2022), T-DXd-dependent TopI poisoning likely increases a demand for TopII α within a window of the cell cycle when TopII α is most abundant, making it a candidate target for additional therapeutic intervention. In support of this rationale, our data demonstrate that TopII targeting with doxorubicin in the T-DXd treatment setting elevates DNA damage, checkpoint pathway activation, and the suppression of tumour spheroid growth (Figures 2 & 3). These data therefore highlighting the potential utility of T-DXd/doxorubicin combination therapy as a therapeutic strategy for the HER2-low setting.

In addition to processing DNA topologies within S phase, TopII α also plays an independent role in the decatenation of entangled chromosomes at the G2/M transition, which is essential for faithful chromosome resolution. During S phase, TopI and TopII α cooperate to dissipate torsional stress that accumulates ahead of the advancing replisome. Fork rotation can also relieve positive torsional stress ahead of the replisome by converting it into sister chromatid intertwines (precatenanes) within the newly replicated daughter duplexes. TopII α acts behind the replication fork to remove these precatenanes, thereby facilitating faithful chromosome resolution later at the G2/M transition (Cebrián et al., 2015; Lucas et al., 2001). TopII α poisoning would be predicted to increases the burden of entangled chromosomes that persist to G2/M, a phenotype that would be expected to be further exacerbated by TopI inhibition, and which places further demand on TopII α-dependent decatenation at the G2/M transition. This likely explains the enhanced cytotoxicity, DNA damage burden and checkpoint pathway activation associated with T-DXd/doxorubicin combination therapy.

While TopII α poisoning directly compromises chromosome decatenation, TopI poisoning primarily perturbs DNA replication and is therefore expected to increase the burden of under-replicated DNA in cells that escape the intra-S checkpoint and progress into G2. PARP plays a primary role in the removal of trapped TopIcc and the resolution of the damaged lesion by coordinating enzyme ubiquitination, proteolytic trimming and excision, and subsequent DNA backbone re-ligation. And this is likely why in our study we see a robust DNA damage signature and a dramatic reduction in cell viability for HER2-low tumour spheroid models in the T-DXd/olaparib combination setting (Figure 2E & F, Figure 3B, E, & F).

In addition to the acute response provided by the PARP/TDP1 pathway in the resolution of trapped TopIcc, MUS81 has also been reported to play an important role in facilitating replication fork progression and promoting cell survival in response to TopI poisons (Regairaz et al., 2011). Regairaz et al. argue that MUS81-dependent cleavage of replication forks provides an alternative mechanism for dissipating excessive supercoiling in the absence of productive topoisomerase activity, allowing DNA replication to progress under conditions of TopI poison-induced replication stress. In our study using HER2-low CRC spheroids, we show that T-DXd promotes far greater cytotoxicity and DNA damage in a MUS81-deficient *vs.* MUS81-wildtype background (Figure 3A-F, Figure 4B & E). Under conditions of T-DXd-dependent genotoxic stress, the S phase functions of MUS81-EME2 are likely of particular importance in sustaining DNA replication through stalled fork processing. However, MUS81 plays two temporally and functionally distinct roles in resolving DNA replication during the cell cycle, both of which are likely to become increasingly important under conditions of T-DXd-dependent replication stress. For those cells that successfully escape S phase, and that likely harbour an increased burden of under replicated DNA, there is likely an increased reliance on the SLX4-MUS81–EME1 pathway that mediates mitotic DNA synthesis (MiDAS) (Pepe & West, 2014). Within this context, it is interesting that heterozygous inactivating mutations in the SLX4 gene have been associated with acquired secondary resistance in the HER2-positive and HER2-low metastatic breast cancer setting (Mosele et al., 2023). Given the sensitizing effects of MUS81 loss to T-DXd that we have observed in our study, this raises the intriguing possibility that haploinsufficiency in SLX4 expression, which is predicted to impact on the G2/M MiDAS functions of MUS81 but not MUS81-EME2-related fork processing in S phase, enables intact MUS81 fork processing while attenuating the rate of SLX4-MUS81-EME1-mediated MiDAS that, with an abundance of under replicated DNA substrates, could otherwise result in chromosomal pulverization if left unchecked. By leaving the SLX4-independent S phase functions of MUS81 unhindered, while maintaining a permissive but attenuated level of MiDAS, SLX4 haploinsufficiency may provide an environment in which a temporal alteration in the efficiency of MUS81 activity is more favourable for survival under conditions of chronic replication stress. Additionally, recent evidence points to a separate role for MUS81 in the resolution of trapped TopIcc in mitosis, via a TDP1-dependent mechanism (Paul Chowdhuri & Das, 2024), suggesting that MUS81 can function both temporally and spatially throughout the cell cycle, by engaging different signalling partners and scaffolds, to counter the genotoxic effects of TopI poisons.

Within G2, TopII α and MUS81 play distinct but complementary roles in chromosome resolution that are essential for the faithful completion of cell division, through the resolution of chromosomal decatenation and MiDAS, respectively. We show that, in the context of T-DXd therapy, perturbation of either of these pathways enhances the DNA damage burden and elevates cytotoxicity in the HER2-low setting, highlighting the potential utility of T-DXd/doxorubicin combination therapy or the application of single-agent T-DXd in the MUS81 low-expression setting. We also show that T-DXd-dependent induction of cellular arrest within S and G2 is coincident with maximal TopII α availability, highlighting this enzyme as both a candidate therapeutic target and potential marker of response to T-DXd therapy.

Collectively, the DESTINY trials have demonstrated the clinical benefit of T-DXd therapy over a broad range of HER2-positive solid tumour types (Meric-Bernstam et al., 2024; Raghav et al., 2024; Smit et al., 2024), and tumour-agnostic factors will likely play an increasing role in clinical decisions governing the administration of this and future ADCs. As the clinical indications that guide therapy decisions become increasingly biomarker led, there is a critical need to broaden our understanding of the molecular signatures that influence treatment response beyond expression of the target antigen alone. In this study, we identify TopII α as a marker of response and a target for combination therapy in the T-DXd treatment setting using HER2-low tumour spheroid models of breast cancer and CRC. Additionally, we demonstrate enhanced T-DXd efficacy in HER2-low CRC spheroids deficient in MUS81 expression.

FDA approval for T-DXd use in metastatic HER2-positive solid tumours highlights the extended clinical application of this ADC to aggressive tumour types beyond the BC setting (F.D.A, 2024). Reduced MUS81 expression has been associated with poorer prognosis in both hepatocellular carcinoma (HCC) and CRC. In CRC, MUS81 expression loss is associated with increased incidence of liver metastasis (Wu et al., 2011). Additionally, suppression of MUS81 expression has been shown to promote sensitivity to camptothecin and epirubicin in tumour xenograft models of Serous Ovarian Cancer (Lu et al., 2019) and HCC (Wu et al., 2016), respectively. Taken together, our findings and those of others suggest the possibility that T-DXd therapy, either as a single agent or in combination with TopII α or PARP inhibitors, may have utility in a HER2-positive/ MUS81-low subgroup of patients, for which prognosis is particularly poor.

We believe that these findings warrant future investigation into the clinical utility of TopII α both as a marker of response and a target for combination therapy, and the potential utility of MUS81 expression status for stratifying patients with HER2-positive disease, in the T-DXd treatment setting.

## MATERIALS AND METHODS

### Spheroid culture and treatment

HCC1954 and T47D BC cells were cultured in RPMI 1640 medium, and DLD1 CRC cells in advanced DMEM (DLD1) supplemented with 10% FBS, 1% L-Glutamine, and 1% penicillin/streptomycin. All cell lines were maintained at 37°C in a stable humidified atmosphere of 5% CO_2_. Cell spheroids were grown in ultra-low adherence round-bottomed 96-well plates (Corning). Cell stocks were trypsinized, dissociated to single cells, and seeded at a density of 3000 cells per well in a final volume of 100 μl. Cells were observed at day 6 to confirm spheroids formation prior to the addition of treatments. At day 3 post-treatment, spheroids were collected using wide-bore, ultra-low protein bind pipette tips and processed for imaging and western blotting.

Stocks of 20.6 mg/mL T-DXd (provided by Daiichi Sankyo), 20 mg/mL T-DM1 and trastuzumab (sourced from clinic), 10 mM doxorubicin (ApexBio, A3966) and 10 mM olaparib (ApexBio, A4154) were diluted in complete medium to the desired final concentrations. When working, precaution was taken to protect all drug stocks and working solutions from light.

### Light-sheet imaging

Spheroids were collected after treatment, washed briefly in PBS and fixed in 4% PFA at room temperature (RT) for 60 min. Samples were then washed in PBS for 30 min twice and transferred into a 24-well plate with a LabelEase insert (Idylle). Samples were incubated in 500 µL 0.5% triton-x for 30 min twice and then blocked for 60 min at RT in 500 µL blocking buffer (0.1% Triton-X100 + 2% BSA + 0.02% Sodium Azide in PBS). Primary antibodies were diluted in 350 µL blocking buffer and incubated overnight at 37 °C. Samples were washed twice with 500 µL 0.2% triton-X100 for 30 min at RT, followed by incubation in 350 µL secondary antibodies in blocking buffer for 60 min at 37 °C. Samples were then washed in 0.2% triton-x for 30 min twice and incubated in 350 µL directly conjugated antibodies and DAPI in blocking buffer overnight at 37 °C. Next, samples were washed twice in 0.2% triton-X100 for 30 min and twice in PBS for 30 min. Samples were placed in disposable base moulds (Simport, M475-1) and embedded in 1% low melting point agarose (16520-050, Invitrogen). Once the agarose set, the agarose sample blocks were placed back into the 24-well plate without the insert. The blocks were then dehydrated by 2x 60 min RT incubations in increasing concentrations of 25%, 50%, 70% ethanol and finally overnight in 100% ethanol. For sample clearing, the agarose blocks were transferred to a polypropylene plate (43001-0066, Ritter) and incubated in 2 mL ethyl cinnamate (A12906.0E, Alfa Aesar) for 2 hr twice and then incubated overnight at 37 °C. At this stage, the samples were stored at RT in the dark until imaging. All steps were carried out in the dark and done on an orbital shaker (70 rpm).

Image acquisition was carried out using an Alpenglow light-sheet microscope, using the LUMI 2.1 software and image processing was carried out using Alpenglow’s 3Dm software. Image analysis was carried out with Imaris Viewer x64 9.9.1.

### Western blotting

Spheroids were collected after treatment, briefly washed with PBS and lysed in 100 µL of RIPA lysis buffer (Thermo Scientific, #89900) + ReadyShield Protease and Phosphatase Inhibitor Cocktail (Sigma-Aldrich, #PPC2020) for 1 hour on ice. The lysates were clarified by centrifugation and transferred to fresh tubes for protein quantification by Pierce BCA Protein Assay (Thermo Scientific, #23225). Equal amounts of protein were loaded onto precast 4–12% gradient Bis-Tris gels (Invitrogen, #NP0335BOX). Gels were run using NuPAGE MOPS SDS running buffer (20X, Invitrogen #NP0001) in an XCell SureLock mini-cell chamber (Invitrogen, #10572913). Proteins were transferred to a 0.45 μm Immobilon-P PVDF membrane (Millipore, #IPVH00010), pre-activated with 100% methanol, using an XCell II blot module (Invitrogen, #10572913). Transfer was performed in ice-cold transfer buffer (10% methanol, 50 mL 20X NuPAGE transfer buffer [Invitrogen #NP00061], 850 mL Milli-Q water).

Following transfer, membranes were washed in 1X TBST (10X TRIS-buffered saline [Thermo Scientific #J60764.K3], diluted with Milli-Q water and 0.05% Tween-20) and blocked with 5% non-fat dry milk (Millipore, #70166) in TBST at RT for 1 hr. The membranes were washed in TBST and then incubated overnight at 4 °C with primary antibodies diluted in 3% BSA (fraction V, Roche #10735108001D2) in TBST. Membranes were washed again and then incubated with HRP-conjugated secondary antibodies (Dako) for 1 hour at room temperature. After a final wash in TBST, immunoreactivity was detected using SuperSignal West Pico Plus chemiluminescent substrate (Thermo Scientific, #34580) using a G-Box imaging system (Syngene, India).

All primary antibodies used in this study are recorded in Supplementary Table 1. HRP-conjugated Goat anti-Rabbit (Dako, P0488, 1:3000) and Goat anti-Mouse (Dako, P0447, 1:3000) IgG were used for secondary detection.

### Cell viability assay

Spheroids were seeded and grown for 3 days before treatment. Brightfield images were acquired at 0, 6, 9, 12 and 15 days post treatment, to enable spheroid growth to be measure over time (mean diameter measurements, ImageJ). At day 15, cell viability was quantified using CellTiter-Glo 3D Cell Viability (Promega, G9683). Briefly, 50 µL of medium was removed from each well and the remaining medium was mixed with 50 µL of the CellTiter-Glo reagent. The plate was then incubated in the dark at RT with shaking (200 RPM, 1 h) before lysate transfer to a white-walled analysis plate (Thermo Scientific #10479501). Luminescence was measured using a CLARIOstar Plus microplate reader (BMG Labtech). Dose-response curves and spheroid mean-diameter graphs were generated using GraphPad Prism 10.4.1 (GraphPad Software Inc., La Jolla, California, USA).

### Flow Cytometry

Organoids were dissociated to a single cell suspension, washed in PBS, and then stained with Zombie UV Fixable Viability Kit (Biolegend, #423108). Subsequently, cells were washed in FACS buffer, fixed with Fluorofix (BioLegend, #422101), then incubated in Intracellular Fixation and Permeabilization buffer (Invitrogen, #88-8824-00), before staining with SYTO61 nuclear dye (Invitrogen, #S11343). Flow cytometry was performed on a CytoFLEX LX flow cytometer (Beckman Coulter, CA, USA) at the Advanced Cytometry Platform, Guy’s Hospital. Data analysis was performed using FlowJo v10 Software (BD Life Sciences).

### Cell line generation

T47D and DLD1 CAS9 stable cell lines were generated by lentiviral mediated gene transduction. Briefly, lentivirus carrying CAS9 and blasticidin S resistance transgenes was packaged in HEK 293T cells using the third-generation lentiCas9-Blast transfer vector (Addgene, #52962) and the pMDLg/pRRE (Addgene, #52962), pRSV-REV (Addgene, #52962), and pMDG.2 (Addgene, #52962) accessory plasmids, with PEI as the transfection reagent. Target cells were transduced in 6-well plates using unconcentrated, 0.22-micron filtered viral supernatant in the presence of 4 µg/mL Polybrene (Santa Cruz, SC134220) for 6 h. After medium replacement, cells were cultured for an additional 48 h before the selection of clones with successfully integrated transgene using blasticidin (4 µg/mL). Cas9/EGFP dual-expressing cells were generated as above by transducing the CAS9 parental lines with lentivirus packaged using the third-generation pLJM1-EGFP transfer vector (Addgene, #19319) that encodes EGFP and confers puromycin resistance (1 µg/mL for selection).

T47D HER2-KO cells were generated by subjecting the T47D CAS9 parental line to two sequential rounds of transient transfection with a mix of two human HER2-directed Edit-R predesigned sgRNAs (Horizon Discovery, SG-003126-01 & SG-003126-02) using DharmaFECT 2 Transfection Reagent (Horizon Discovery, T-2002). The T7 Endonuclease assay was used to confirm a high efficiency of gene editing before performing a serial dilution of the cell suspension diagonally across 96-well culture plates (8,000 cell starting density) for the derivation of clonal cell populations. After two weeks culture, wells confirmed to contain single colonies were selected and the cells expanded. Loss of HER2 protein expression was evaluated by western blotting and three confirmed HER2-negative clones then selected and pooled to generate the experimental T47D HER2-KO cell line. DLD1 MUS81-KO cells were generated following the same method, using a mix of two human *MUS81*-directed Edit-R predesigned sgRNAs (Horizon Discovery, SG-016143-01 & SG-016143-02). Three MUS81-negative clones that exhibited comparable HER2 expression to the parental line (to ensure equality in ADC target antigen expression) were then selected and pooled to derive the DLD1 MUS81-KO experimental line.

### Spheroid cryosectioning and immunoffuorescence analysis

At the end of drug treatment assays, spheroids cultured in ultra-low adherence U-bottom 96-well plates were harvested and all spheroids for a given treatment group pooled together before fixation, processing for OCT embedding, cryosectioning, and subsequent immunostaining and imaging by epifluorescence wide-field microscopy. In imaging assays, where direct comparisons were made between wildtype (EGFP-positive) and knockout (EGFP-negative) cell lines, the corresponding treatment groups for each line were pooled after fixation. This produced mixed populations of wild-type and knockout spheroids for each given treatment group, with the origin of each spheroid determinable by its EGFP expression status.

Spheroid processing was as follows: Individual spheroids were harvested from 96-well plate wells using low-protein bind wide-bore filtered tips and then pooled in 15 mL Falcon tubes according to treatment group. Spheroids were allowed to settle by gravity before the supernatant was carefully removed and discarded. The spheroids were then washed gently in PBS and fixed in 4% PFA for 1 h. Post-fixation, spheroids were washed 2x in PBS and then incubated over-night in 30% sucrose/PBS (w/v). The following day, the spheroids were transferred to a 1:1 mix of 30% sucrose/PBS:OCT and incubated for 1 h at RT with rolling. The spheroids were then gently pelleted by centrifugation at 200 × g for 5 min, the supernatant discarded, and the pellet then resuspended in 100% OCT with gentle mixing using a wide-bore pipette tip. After a 1 h incubation at RT, the spheroids were pelleted again by centrifugation and then transferred to cryomoulds (Simport, M475-1). Moulds were briefly centrifuged at 200 x g for 2 min to ensure all spheroids collected at the base, and the sample then frozen on powdered dry ice before storing at -80°C. OCT blocks were sectioned at 4.5-micron thickness using a Leica cryostat. Sections from all treatment groups within a given experiments were arrayed together on a single slide. This enabled immunolabelling and imaging of spheroids from all treatment groups to be conducted together on the same slide, to facilitate statistical analysis of biomarker expression when performing inter-group comparisons, as well as intra-group comparisons in cases where control/EGFP-positive and knock-out/EGFP-negative spheroid models were used.

Slide staining was performed according to standard immunofluorescence protocols. Slides were then imaged by tile-scanning using a Nikon Eclipse Ti-2 inverted wide-field epifluorescence microscope equipped with a 20x ELWD objective, Nikon DS-Qi2 sCMOS camera, motorized stage, and NIS-Elements acquisition software. Post acquisition image analysis was performed in ImageJ using purpose written macros for image feature extraction, which included whole-spheroid and EGFP subpopulation masking, and subsequent nuclei masking for the compartmentalised analysis of marker fluorescence intensity values. All image processing and analysis macros and an explanation of their can be made available upon request.

### Statistical analysis

All statistical analyses and graphical representations of data were performed and generated using Prism 10 software (GraphPad, V10.6.0). One-way and two-way ANOVA with multiple comparisons were used to test the significance of differences between treatment groups. In all cases, asterix were used to indicate degree of significance as: *, p < 0.05; **, p < 0.01; ***, p < 0.001.

## ACKNOWLEDGEMENTS

This research was funded by the KCL/GSTT Centre for Translational Medicine The authors declare that they have no conflict of interest.

---

**Figure S1:**
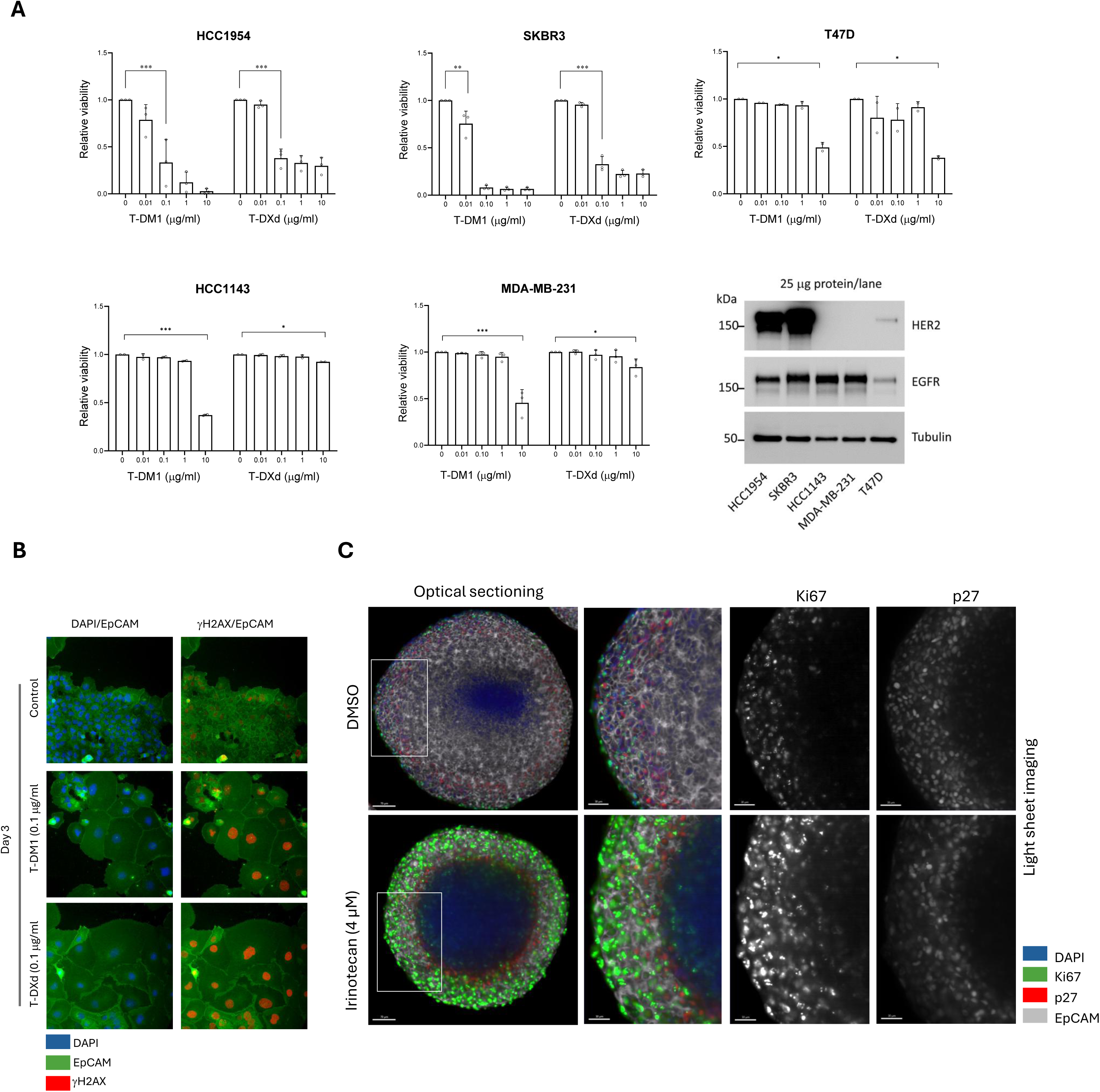
HER2 expression dictates T-DXd efficacy in *in vitro* models of breast cancer. (A) Alamar blue-based cell viability analysis of breast cancer cell lines of varying HER2 expression subjected to the indicated concentrations of T-DM1 and T-DXd within the two-dimensional *in vitro* culture setting. Bars indicate viability relative to untreated controls. The associated western blot shows HER2 expression levels for the different breast cancer cell lines studied. (bar graphs: N = 2 repeats, T47D; N = 3, all other lines). (B) Immunofluorescence analysis of γH2AX expression and cell morphology demonstrate prevalent ADC-dependent DNA damage and cell enlargement in response to both T-DM1 and T-DXd. (C) Whole-organoid light-sheet microscopy analysis of the anatomical organisation of the cancer spheroid reveals concentric zones of proliferation, quiescence and necrosis that are disrupted by TopI poisoning.

**Figure S2:**
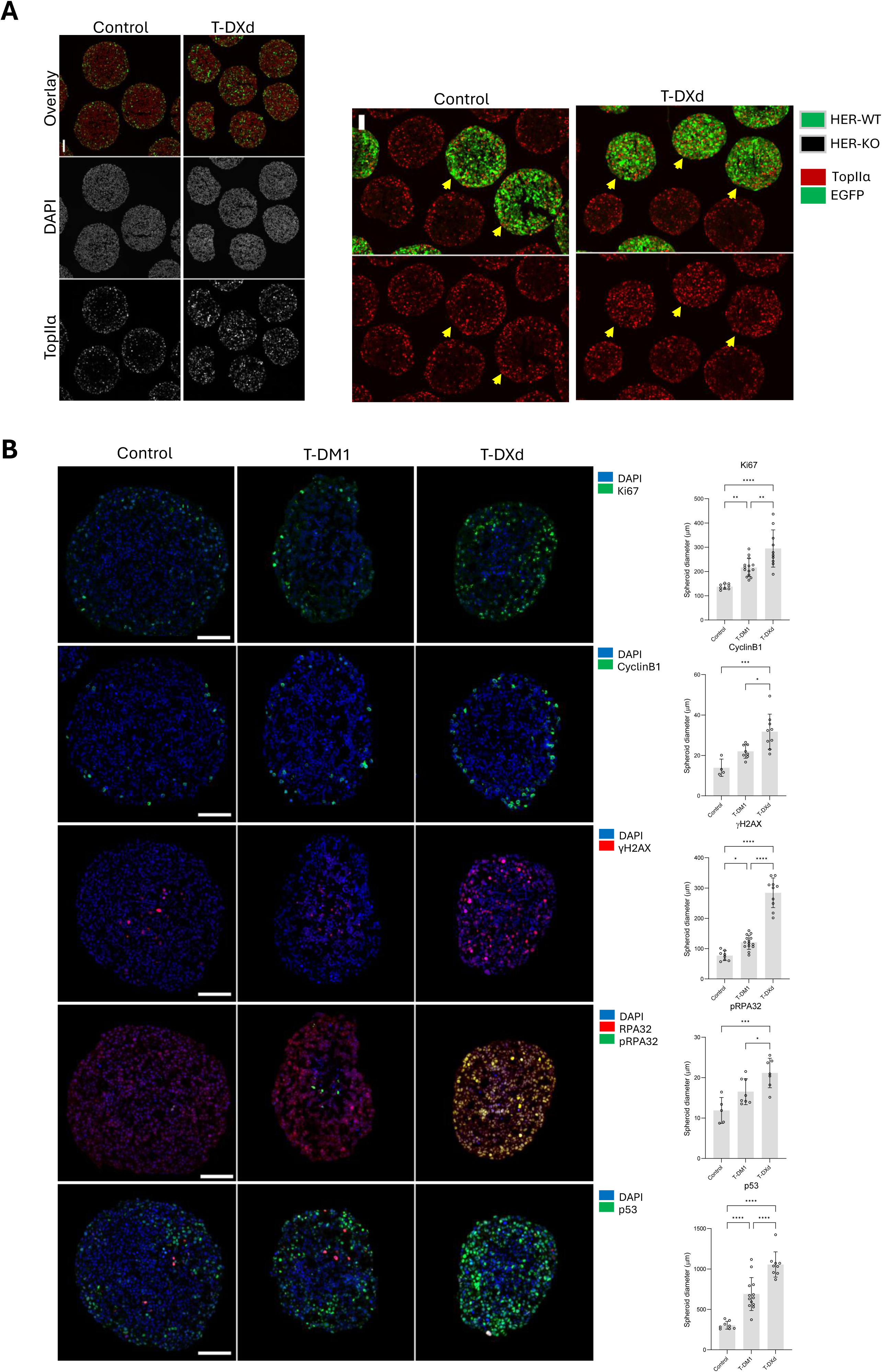
T-DXd promotes markers of DNA damage and proliferation in HER2-low breast cancer spheroids. (A) Left hand panel: Cryosections of control and T-DXd-treated T47D HER2-low spheroids showing therapy induced induction of TopII α expression. Right-hand panel: Syngeneic HER2-KO (EGFP-negative) T47D spheroids were included in T-DXd experiments to provide a base-line control for potential off-target effects of drug in the HER2-low setting. Yellow arrows point to corresponding spheroids in the upper and lower rows. Top row: TopII α staining + EGFP overlay, bottom panel, TopII α staining alone. (B) Representative immunolabelled cryosections of T47D spheroids from control, T-DM1 and T-DXd stained for markers of DNA damage, replication stress, and proliferation. Right hand graphs: Quantitative analysis of per-spheroid mean intensities for the corresponding markers in the images on the left. Each data point represents the mean intensity score for a single spheroid (N≥4 independent spheroids per treatment group).

**Figure S3:**
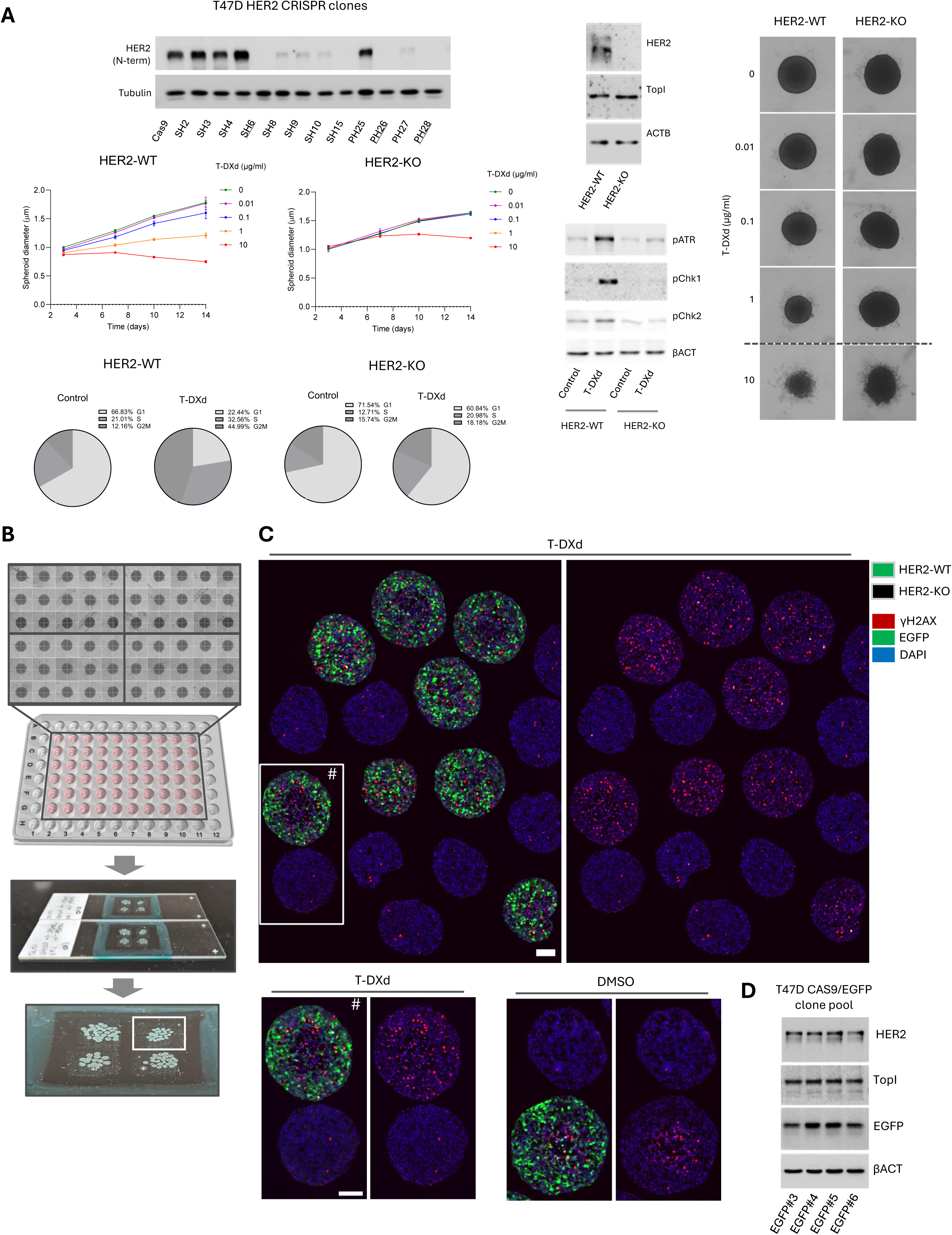
Imaging-based assay for the analysis of mixed populations of CRISPR-engineered tumour spheroids. T47D HER2-KO CRISPR models were generated to discriminate between the on-target and off-target effects of ADC under our experimental conditions. (A) A T47D HER2-KO cell line was generated by pooling three different clones generated from CRIPSR-mediated gene silencing. Western blotting demonstrates HER2 expression across a range of T47D clones generated during the CRISPR HER2 knock-out selection process. The three clones subsequently pooled for the HER2-KO cell line are underlined. Expression of HER2 (T-DXd antigen target) and TopI (T-DXd payload target) between the control and HER2-KO lines is shown in the western blot on the right. Spheroid growth assays (middle left panel) and cell cycle analysis (bottom left panel) using the syngeneic HER2-WT and HER2-KO lines demonstrate the specificity of the T-DXd response to presence of the ADC target antigen. Western blotting (bottom right) demonstrates that HER2-KO induced T-DXd resistance is associated with a concomitant loss in DNA-damage response checkpoint pathway activation. The far-right image panel shows representative brightfield images of T47D HER2-WT (left column) and HER2-KO (right column) spheroids cultured for 10 days in the indicated concentrations of T-DXd. (B) CRISPR-based syngeneic tumour spheroid assay development. Tumour spheroids were initiated using ultra-low adherence U-bottom 96-well plates, which enable the highly reproducible generation of uniformly sized cancer spheroids in individual wells. Upper image: montage of phase contrast images of all tumour spheroids from the central 60 wells of the 96-well plate, demonstrating the highly uniform size and morphology of the cultures. Middle image: representation of a 96-well plate to show the culture arrangement. Lower images: photograph of a glass slide with cryosections of tumour spheroids from four treatment groups arranged as a 2 x 2 matrix. Up to six treatment groups were arranged in this way. Each treatment group includes both WT and target gene-KO spheroids, with the WT line (CAS9 control) engineered to express EGFP to enable discrimination between the two spheroid populations. (C) Epifluorescence images of HER2-WT (EGFP-positive) and HER2-KO (EGFP-negative) spheroids imaged together within the same field of view. This imaging assay workflow allowed for the immunolabelling and imaging of spheroids from all treatment groups to be imaged together on the same slide, to facilitate statistical analysis of biomarker expression when performing inter-group comparisons. (D) Western blot demonstrating uniform EGFP, HER2 and TopI expression among four T47D CAS9/EGFP cell clones that were pooled to derive the control line.

**Figure S4.**
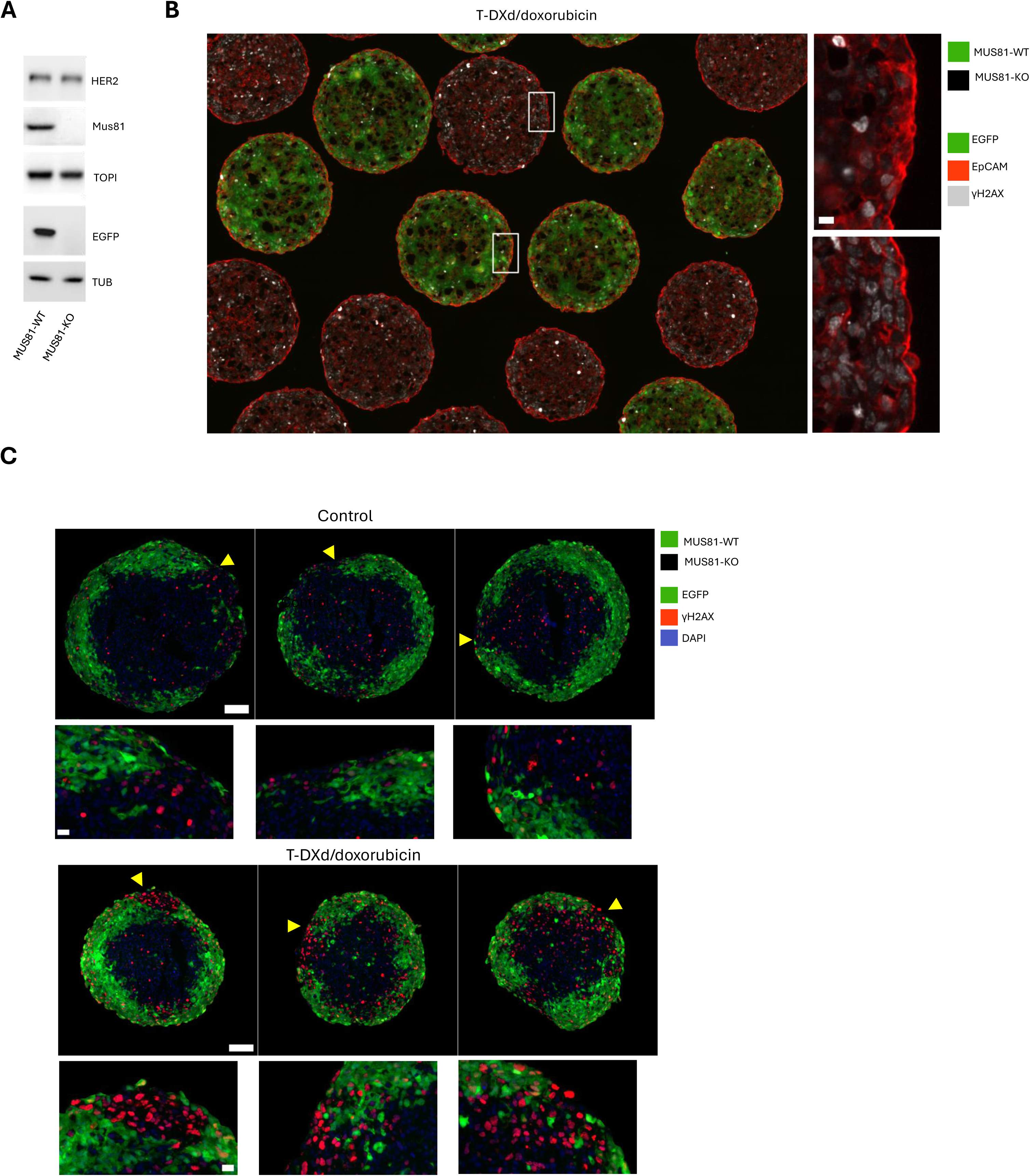
MUS81 loss sensitises HER2-low CRC spheroids to T-DXd/doxorubicin combination therapy. (A) Western blot demonstrating relative expression of MUS81, HER2 & TOPI between the control MUS81- WT/EGFP-positive and MUS81-KO/EGFP-negative syngeneic DLD1 cell lines used in mono and chimeric spheroid studies. (B) Comparative immunofluorescence analysis of the γH2AX DNA damage marker in a mixed population of EGFP MUS81-WT and MUS81-KO DLD1 spheroids following challenge with T-DXd/doxorubicin combination therapy. The associated zoom regions highlight the elevated DNA damage within the periphery of MUS81-deficient spheroids, when compared with wild-type (EGFP-positive) spheroids. (C) Immunofluorescence analysis of chimeric MUS81-WT/MUS81-KO DLD1 spheroid sections from control (upper) and T-DXd/doxorubicin treatment groups, demonstrating elevated and extensive DNA damage within the MUS81-KO population of the combination treatment group. Arrows highlight examples of intense γH2AX nuclear staining in MUS81-KO cells (EGFP-negative).

**Supplementary table 1.** List of all antibodies used in this study and their associated applications.

| **Marker** | **Clone** | **Company** | **Catalogue number** | **Dilution** | **IgG** | **Use** |
| --- | --- | --- | --- | --- | --- | --- |
| EpCAM Alexa Fluor 647 | EPR20532-225 | Abcam | ab237396 | 1:100 | Rabbit | AG/IF |
| γ-H2A.X (S139) Alexa Fluor 555 | EP854(2)Y | Abcam | ab206900 | 1:200 | Rabbit | AG/IF |
| Ki67 Alexa Fluor 488 | EPR3610 | Abcam | ab197234 | 1:100 | Rabbit | AG/IF |
| p27 Kip1 | D69C12 | Cell Signaling | 3686 | 1:100 | Rabbit | AG/IF |
| CyclinB1 Alexa Fluor 555 | Y106 | Abcam | ab214381 | 1:100 | Rabbit | IF |
| Human IgG | IG26 | Abcam | ab200699 | 1:1000 | Mouse | IF |
| DXd | YA897 | MeedChemExpress | HY-P81054 | 1:1000 | Mouse | IF |
| HER2 | 29D8 | Cell Signaling | 216S | 1:1000 | Rabbit | WB/IF |
| TopII $\alpha$ | D10G9 | Cell Signaling | 12286S | 1:1000 | | WB/IF |
| TopI | Polyclonal | Cell Signaling | 79971S | 1:1000 | Rabbit | WB/IF |
| pChk1 (Ser345) | 133D3 | Cell Signaling | 2348 | 1:1000 | Rabbit | WB/IF |
| pChk2 (T68) | C13C1 | Cell Signaling | 2197 | 1:1000 | Rabbit | WB/IF |
| P53 | DO-1 | Santa Cruz | Sc-126 | 1:1000 | Mouse | IF |
| RPA32 | EPR2877Y | Abcam | ab76420 | 1:200 | Rabbit | WB/IF |
| MUS81 | MTA30<br>2G10/3 | Abcam | ab14387 | 1:500 | Mouse | WB |
| γ-H2A.X (S139) | 20E3 | Cell Signaling | 9718 | 1:1000 | Rabbit | WB |
| GAPDH | 1E6D9 | Proteintech | 60004-1-Ig | 1:1000 | Mouse | WB |

IF: Immunofluorescence microscopy
AG: Alpenglow light-sheet imaging (where not included above)
WB: Wester blotting

